# Mutation experiments with AcrB and MexB D408A mutants indicate the efflux liability of antibiotics

**DOI:** 10.1101/2025.01.06.631490

**Authors:** Maria Laura Ciusa, Vito Ricci, Laura JV Piddock

## Abstract

**Background:** We showed that exposure of an AcrB D408A mutant to efflux inhibitors applied evolutionary pressure to select bacteria with the wild type *acrB* sequence. This suggested that reversion to wild type can differentiate between efflux inhibitors. Thus, we hypothesized that this experiment could identify inhibitors of the primary RND pump, AcrB or its homologues in other species.

**Objectives:** To construct three mutants, *Escherichia coli* AcrB D408A, *Klebsiella pneumoniae* AcrB D408A and *Pseudomonas aeruginosa* MexB D408A and expose the mutants to substrates and non-substrates of AcrB and/or MexB and determine the rate of reversion to wildtype *acrB/mexB* sequence.

**Methods:** Mutant *Escherichia coli* AcrB D408A, *Klebsiella pneumoniae* AcrB D408A and *Pseudomonas aeruginosa* MexB D408A were constructed with site-directed mutagenesis of the relevant nucleotide in the *acrB/mexB* gene. Mutants were exposed on agar to substrates and the mutation frequency and mutation rate determined. The MIC of antibiotics and the presence/absence of the D408A substitution was determined for mutants.

**Results:** Exposure to the AcrB substrates chlorpromazine and minocycline reverted the D408A genotypes to wild type in a species-dependent manner. Exposure to a non-AcrB substrate, spectinomycin, did not select wild type *acrB*. Chlorpromazine selected for wild type *acrB K. pneumoniae* as it had for *S*. Typhimurium, whereas minocycline selected for wild type *E. coli acrB*. None of the antibiotics selected wild type *mexB*, including the tested MexB substrates.

**Conclusions:** Evolutionary paths depend upon the genetic background of the species and availability of alternative routes/genetic pathways that can confer resistance/decreased susceptibility to antibiotics.

## Introduction

The WHO global priority pathogens list for which there is a critical need and high priority for new antibiotics is dominated by Gram-negative bacterial species.^1^ It is easy to find compounds with antibacterial activity for Gram-positive bacteria, but it remains difficult to find those effective against Gram-negative bacterial species. This scientific challenge is due to two factors (1) a permeability barrier, the outer membrane, that prevents access of many antibiotics to their intracellular target, and (2) the presence of multidrug resistance (MDR) tripartite efflux pumps that export many compounds from the bacterial cell. ^2^ There is a natural synergy between these two mechanisms. ^2, 3^ Under laboratory conditions, gene deletion, mutational inactivation, introduction of a loss of function mutation in a gene encoding an RND type MDR efflux pump such as AcrB, or inhibition of MDR efflux systems causes increased susceptibility to a wide range of structurally dissimilar antibiotics. ^2, 4^ For many species, there is also loss of virulence, ability to form a biofilm, and the frequency of resistance to antibiotics is markedly reduced. ^2^ For many years, studies that have carried out screening to identify new antibacterial compounds have utilised mutants in which the gene coding for an efflux pump component (e.g., *Escherichia coli acrB, E. coli tolC* or *Pseudomonas aeruginosa mexB*) have been inactivated or deleted. ^2^

Efflux pump proteins such as AcrB and MexB have been studied extensively and have been shown to be proton motive force-dependent and function via a drug/proton antiport mechanism. *E. coli* AcrB and *P. aeruginosa* MexB contain two acidic residues, Asp408 and Asp407, in the transmembrane (TM) helix 4 that have been shown to be essential for transport function ^5, 6^. For many years it has been assumed that the phenotype resulting from inactivating or deleting the efflux pump genes equates with loss of efflux. However, until 2017 it was unclear as to whether the phenotype of multidrug susceptibility was due to the loss of efflux function rather than the lack of a large integral membrane protein. ^4^ Wang-Kan et al., explored this question by constructing a mutant *Salmonella enterica* serovar Typhimurium with a single amino acid substitution in AcrB (D408A). ^4^ The mutant was able to produce its AcrB protein however that mutant protein could not derive energy from the PMF because it could not translocate protons and therefore could no longer actively efflux substrates. The ‘loss of AcrB efflux function’ D408A mutant was hyper-susceptible to antibiotics, dyes and disinfectants, the same as an isogenic *acrB* deletion mutant. Both *acrB* deleted and D408A mutants accumulated much higher levels of fluorescent dyes and antibiotics. Both mutants were also less able to infect tissue culture cells and mice. However, in the absence of AcrB over half of the genes in the *Salmonella* genome had altered expression, but far fewer genes were altered when there was loss of efflux in the AcrB D408A mutant. This shows that much of the altered transcriptome was due to loss of a large membrane protein and not loss of efflux function. Furthermore, previous studies have shown that when the AcrB protein is lacking, there is a concomitant increase in production of homologous RND efflux pumps such as AcrD and AcrF ^7, 8^. In the D408A mutant these proteins were not overproduced. Indeed, in the D408A mutant AcrD and AcrF were produced at virtually undetectable levels, which means that they cannot compensate for the loss of AcrB efflux function. ^4^ This is important because strategies to discover efflux pump inhibitors have assumed that inhibition of AcrB, AcrD and AcrF will be required; this may not be necessary.

In another previous study, ^9^ we showed that exposure of a *S*. Typhimurium AcrB D408A mutant to AcrB substrates and competitive efflux inhibitors such as chlorpromazine and amitriptyline applied evolutionary pressure to select bacteria with the wild type *acrB* sequence. These efflux inhibitors selected for reversion of the D408A sequence to wild type *acrB* more often than the AcrB substrates ethidium bromide or minocycline. When the D408A mutant was exposed to the non-AcrB substrate spectinomycin, AcrB D408 was retained in the resulting antibiotic-resistant mutants. Therefore, the reversion of the AcrB D408A to wild type appears to differentiate between compounds with efflux inhibitory properties, suggesting there is the potential for this experiment to be used to identify inhibitors of the primary RND pump, AcrB or its homologues in other species.

To test our hypothesis, in the present study we constructed three more mutants, *E. coli* AcrB D408A, *Klebsiella pneumoniae* AcrB D408A and *P. aeruginosa* MexB D408A and exposed them to substrates and non-substrates of AcrB and/or MexB and determined the rate of reversion to the wild type sequence. Data obtained were compared with those for the *S*. Typhimurium AcrB D408A mutant. ^9^

## Methods

### Strains, media, and construction of the *K. pneumoniae* AcrB D408A and *P. aeruginosa* MexB D408A mutants

All strains used in this study are shown in Table 1. Bacterial strains were grown overnight at 37°C in Lennox broth (Sigma-Aldrich, UK) or Iso-Sensitest broth (Oxoid, UK). All chemicals and antibiotics were supplied by Sigma-Aldrich, UK.

**Table 1.** Frequency and rate of mutation of the D408A mutant after exposure to selecting agents.

| Strain | Selecting drug | Selecting concentration (mg/L) | Mutation frequency (CFU/ml) | Mutation rate (CFU/ml) | Total number of mutant colonies | Percent of colonies with WT *acrB* |
| --- | --- | --- | --- | --- | --- | --- |
| *S. Typhimurium* SL1344 | CPZ | 60 | $1.82 \times 10^{-10}$ | $5.94 \times 10^{-10}$ | 152 | 100 |
| | MIN | 0.5 | $6.48 \times 10^{-10}$ | $1.399 \times 10^{-9}$ | 702 | 2 |
| | SPEC | 128 | $5.46 \times 10^{-10}$ | $1.34 \times 10^{-9}$ | 459 | 0 |
| *E. coli* MG1655 | CPZ | 60 | $9.29 \times 10^{-9}$ | $1.10 \times 10^{-9}$ | 2137 | 4 |
| | MIN | 1 | $1.79 \times 10^{-9}$ | $8.8 \times 10^{-10}$ | 233 | 76 |
| | SPEC | 20 | $3.88 \times 10^{-10}$ | $1.14 \times 10^{-10}$ | 916 | 0 |
| *K. pneumoniae* Ecl8 | CPZ | 64 | $3.63 \times 10^{-9}$ | $5.55 \times 10^{-10}$ | 3275 | 90 |
| | MIN | 2 | $5.4 \times 10^{-11}$ | $1.8 \times 10^{-11}$ | 618 | 0 |
| | SPEC | 40 | $5.49 \times 10^{-11}$ | $3.68 \times 10^{-10}$ | 687 | 0 |
| *P. aeruginosa* PAO1 | ATM | 2 | $2.89 \times 10^{-10}$ | $3.75 \times 10^{-10}$ | 168 | 0 |
| | MEM | 0.25 | $9.49 \times 10^{-10}$ | $2.89 \times 10^{-10}$ | 655 | 0 |
| | IMI | 10 | $1.10 \times 10^{-9}$ | $4.23 \times 10^{-10}$ | 796 | 0 |

CPZ, chlorpromazine; MIN, minocycline; SPEC, spectinomycin; ATM, aztreonam; MEM, meropenem; IMI, imipenem. WT, wild type.

The same method used for the construction of the chromosomal *E. coli* AcrB D408A mutant by Wang-Kan et al., ^4^ was used to construct the *K. pneumoniae* Ecl8 mutant with the primers described in Supplementary Table 1. The inserts were synthesised by GenScript, PCR-amplified from the supplied plasmids using Q5 polymerase (NEB) and purified using a gel extraction kit (Neo BioTech) according to manufacturer’s instructions. The presence of the mutation was confirmed by PCR from colony lysates followed by DNA sequencing using the check primers listed in Supplementary Table 1. PCR conditions were as follows: initial denaturation at 95 °C for 3 min, followed by 30 cycles of 95 °C for 30 s, 55 °C for 30 s, and 72°C for 1 min, and a final extension at 72 °C for 5 mins. PCR amplimers were DNA sequenced to confirm the presence of the desired mutation.

The chromosomal *P. aeruginosa* PAO1 MexB D408A mutant was constructed by a modified version of the method described previously by Hmelo et al.,^10^. Flanking primers were designed upstream and downstream of the *mexB*, sequence. The flanking primers are described in Supplementary Table 2. Primers were also designed to introduce the specific mutation into the *mexB* gene; these are described in Supplementary Table 2. PCR was carried out to produce an upstream and downstream fragment, using a combination of the flanking primer and mutagenic primer as follows: pEX18-*mexB*-Fw + *mexB* (D408A)-Rv and pEX18-*mexB*-Rv + *mexB* (D408A)–Fw. The following conditions were used for all PCR reactions: initial denaturation at 98 °C for 3 min, followed by 30 cycles of 98 °C for 10 s, 58 °C for 30 s, and 72 °C for 15 s, and a final extension at 72 °C for 1 min. The PCR products were analyzed by DNA agarose electrophoresis and purified by QIAquick spin-column (QIAGEN). The pEX18Amp suicide plasmid was linearized by PCR using the following primers: pEX18_fwd 5’–AGCTTGCATGAGCTCGAA TTCGTAATCATGG -3’ and pEX18_rev 5’– AATTCGAGCTCATGCAAGCTTGGCACTG -3’ and the following PCR conditions; initial denaturation at 98 °C for 3 min, followed by 30 cycles of 98 °C for 10 s, 60 °C for 30 s, and 72 °C for 7 mins, and a final extension at 72 °C for 10 mins. Following PCR, DpnI digestion was carried out followed by PCR purification. Gibson assembly was then carried out using the NEBuilder® HiFi DNA Assembly master mix (NEB, cat. no. E5520S). Briefly, the reaction mix contained 10 nM of each amplified upstream and downstream genomic DNA fragment, 5 nM of linearized pEX18Amp plasmid and 10 uL of Gibson assembly master mix. The reaction mixture was incubated at 50 °C for 1 hr and the resulting ligated plasmid was transformed into chemically competent *E. coli* DH5α (NEB). Transformants were checked by colony PCR using the universal pEX18 vector primers, which flank the multiple cloning site: forward, 5’ – GGCTCGTATGTTGTGTGGAATTGTG-3’ and reverse, 5’-GGA TGTGCTGCAAGGCGATTAAG-3’ (annealing temperature: 55 °C). Positive transformants were verified by DNA sequencing. One positive transformant was then used to extract the plasmid and carry out electroporation with competent *P. aeruginosa* PA01 cells. Following electroporation and recovery in LB broth at 37 °C for 2 hrs, cells were plated out onto LB agar containing carbenicillin at 300 µg/mL and incubated at 37 °C for up to 72 hrs. Colonies growing on the LB Carb300 plates were purified by streaking on new LB Carb300 plates. Purified colonies were then streaked onto Tryptone yeast extract plus sucrose agar (TYS10) plates and incubated at room temperature (∼ 23 °C) for up to 7 days. Colonies that grew on the TSY10 agar plates were streaked out onto LB and LB Carb300 plates. Colonies that had lost the integrated plasmid grew on the LB agar plates and not the LB Carb300 plates. Colonies from the LB agar plates were examined by colony PCR using a set of primers flanking the genomic sequence of *mexB* near to the mutation site (Supplementary Table 2). PCR conditions were as follows: initial denaturation at 98 °C for 3 min, followed by 30 cycles of 98 °C for 10 s, 60 °C for 30 s, and 72 °C for 5 mins, and a final extension at 72 °C for 10 mins. PCR amplimers were DNA sequenced to confirm the presence of the desired mutation.

**Table 2.** MICs and *acrB* genotypes of D408A mutants that arose after selective pressure.

| Selecting agent | MIC (mg/L) |  | Mutant selecting concentration | no. of colonies tested | MIC (mg/L) |  |
| --- | --- | --- | --- | --- | --- | --- |
| | Wild type strain | D408A mutant | | | Colonies with wild type *acrB* | Colonies containing AcrB D408A and inhibited by MICs ≥ than those for the wild type |
| *E. coli* MG1655 |  |  |  |  |  |  |
| CPZ | 128 | 32 | 60 | 90/2137 | 256 (**3**) | 256 (**50**)<sup>a</sup> |
| MIN | 2 | 0.5 | 1 | 54/233 | 4 (**41**), 8 (**2**) | 4 (**4**), 8 (**2**)<sup>b</sup> |
| *K. pneumoniae* Ecl8 |  |  |  |  |  |  |
| CPZ | 128 | 32 | 64 | 1/131 | 64 (**2**), 128 (**8**), 256 | 64 (**3**) |
| MIN | 1 | 0.12 | 2 | 15/103 | 0 | 0.5 (**4**), 1 (**80**), 2 (**6**) |
| *P. aeruginosa* PAO1 |  |  |  |  |  |  |
| ATM | 4 | 0.25 | 0.25 | 15/28 | 0 | 4 (**10**), 8 (**80**) |
| MEM | 0.5 | 0.12 | 2 | 90/655 | 0 | 0.5 (**16**), 1 (**74**) |

Number of mutant colonies with each MIC value shown in bold font in brackets. CPZ, chlorpromazine; MIN, minocycline. WT, wild type.
a, 37 tested colonies had the same susceptibility to chlorpromazine as the D408A mutant.
b, Four colonies had the same susceptibility to minocycline as the D408A mutant.

#### Antimicrobial susceptibility

The MICs of antibiotics were determined according to the European Committee on Antimicrobial Susceptibility Testing (EUCAST) methodology using the doubling dilution method on agar. ^11^ As recommended by EUCAST, *E. coli* ATCC 25922 and *P. aeruginosa* ATCC 27853 were used as control strains. Antibiotics were prepared according to the manufacturer’s instructions. The MIC of each agent was recorded as the lowest concentration of compound that prevented visible growth.

#### Mutant selection

An overnight culture (50uL) of each strain of interest was plated onto the surface of Lennox agar supplemented with the selecting agent at the appropriate concentration. To determine which antibiotic and the concentration to use in the mutant selection experiments the MICs of numerous compounds for the D408A mutants and their wild type parental strains were determined on multiple occasions (Table 1 and 2). As a result, *E. coli* MG1655 AcrB D408A and *K. pneumoniae* Ecl8 AcrB D408A were exposed to the *E. coli* AcrB substrates chlorpromazine and minocycline. *P. aeruginosa* PAO1 MexB D408A was exposed to the MexB substrates aztreonam and meropenem. As negative controls, spectinomycin was used as a non-substrate of AcrB for *E. coli* and *K. pneumoniae* and imipenem for *P. aeruginosa*. Mutants were selected by exposure to drug concentrations one or two times the MIC of the antibiotic for the AcrB D408A or MexB D408 mutant strains (Table 1 and 2).

Plates were incubated aerobically at 37°C and checked daily for up to seven days, or until the appearance of colonies, which were then counted. To calculate the CFU/ml in the overnight culture, the culture was diluted 1:1000, followed by three 1:10 serial dilutions in LB broth.

Colonies were replica plated onto fresh Lennox agar with no antibiotic and MacConkey agar to check colonial morphology was consistent with that of the parental strain. Up to 100 mutant colonies per strain/compound combination were collected, cultured in Lennox broth plus 10% glycerol and frozen at -80°C for further analysis.

#### Calculation of Mutation Frequency and Mutation Rate

Consistent with our previous work, ^9^ experiments were done with fluctuation analysis. ^12, 13^ The mutation frequency is the measurement of all the mutants in a given population, while the mutation rate is an estimate of the probability of a mutation occurring per cell per generation. ^13^ The mutation rate was calculated using the MSS maximum likelihood method. ^13^ To ensure confidence in the data from the experiments, agar plates were inoculated with 100 parallel cultures (derived from three biological replicates).

#### Allele-specific qualitative PCR

From each mutant selection experiment, 100 colonies were replica plated onto Lennox agar from the frozen glycerol stocks; all mutants that grew on thawing and sub-culture were screened for the presence of AcrB D408A. To detect reversion of the *acrB* or *mexB* sequences to the wild type, an allelic real time PCR assay was used. Samples were prepared as described by Grimsey et al. ^9^. Primer sequences are listed in supplementary Table 2; the real time PCR reactions were performed on a QuantStudio Real Time PCR instrument (Thermo Fisher Scientific, UK) using the cycling conditions listed in Supplementary Table 2. A melting curve was generated using a temperature range from 50°C to 95°C with increments of 0.5°C every 5 s.

#### Whole-genome sequencing

Representative mutants that retained the original *acrB* D408A genotype and had decreased susceptibility or were resistant to the selecting agent were sent to MicrobesNG (http://www.microbesng.uk) for whole genome sequencing. Paired-end sequencing was carried out on the Illumina HiSeq platform. The reads were mapped to the wild type parental strain reference genome using Bowildtypeie2. Single nucleotide polymorphism (SNP) analysis was carried out with Mpileup, and single nucleotide polymorphisms (SNPs) identified using Artemis (Sanger Institute, United Kingdom).

## Results and Discussion

Exposure to the AcrB substrates chlorpromazine and minocycline reverted the D408A genotypes to wild type in a species-dependent manner (Table 1). Exposure to a non-AcrB substrate, spectinomycin, did not give colonies with the wild type genotype.

After exposure of the *E. coli* and *K. pneumoniae* D408A mutants to chlorpromazine and minocycline the colonies that had a wild type *acrB* genotype also had the same susceptibility as the wild type strains (Table 1). This indicates the evolutionary pressure of these agents was to revert the single nucleotide polymorphism (SNP) in *acrB* to wild type. The mutation frequency and mutation rate were calculated for all drug exposure experiments (Table 1). As found for *S*. Typhimurium, ^9^ the selective pressure of chlorpromazine on the AcrB D408A mutant reverted *K. pneumoniae* to the *acrB* wild type DNA sequence at a high rate (90%) while *E. coli* reverted at a low rate (4%). This suggests that chlorpromazine is a substrate of AcrB in *K. pneumoniae*. The exposure of minocycline also gave different results for *E. coli* versus *S*. Typhimurium and *K. pneumoniae*; with a higher rate of reversion to wild type *acrB* (76%) than the other two species. This suggests that minocycline is a substrate of *E. coli* AcrB and this supports data by others using biochemical methods. ^14^

We also observed that after exposure of the *E. coli* and *K. pneumoniae* D408A mutants to chlorpromazine and minocycline some colonies retained the original D408A genotype and either had similar susceptibility to the wildtype parental strain or required a higher concentration of antibiotic for growth inhibition (Table 2). Despite obtaining increased MICs on several occasions for the mutants selected in the *E. coli* AcrB D408a background, repeated whole genome sequencing (WGS) revealed no single nucleotide polymorphisms (SNPs), inserts or deletions. Indeed, the only SNPs found were in *rhsC*, a rearrangement hotspot protein; ^15^ such proteins have been shown to mediate intracellular competition but have not been previously associated with or implicated in antibiotic resistance. WGS of the *K. pneumoniae* AcrB D408A mutant surprisingly revealed two additional mutations, one in *bcr*, ^16^ the other in *ompR*. ^17^ Bcr is a component in the Bcr/CflA family of MFS efflux transporters and has been previously associated with antibiotic-resistance. ^18^ In *E. coli*, OmpR is part of the EnvZ/OmpR two-component regulatory system involved in regulating the expression of OmpF and in *K. pneumoniae* shown to be involved in virulence. ^17, 19^ WGS of a representative mutant *K. pneumoniae* selected after exposure to minocycline, revealed a SNP in *betI*. ^20^ This is a transcriptional regulator involved in the osmoregulatory choline-glycine betaine pathway of *E. coli* and shown to be upregulated when *K. pneumoniae* is grown as a biofilm. ^21^

Due to high MIC values, we could not use chlorpromazine or minocycline in the experiments with *P. aeruginosa*. We tested spectinomycin and no mutants were selected. We determined the MICs of MexB substrate antibiotics as indicated in the literature ^22^ and chose aztreonam and meropenem for our experiments. However, exposure of *P. aeruginosa* to these agents did not give rise to colonies with the wild type *mexB* DNA sequence. WGS of representative*P. aeruginosa* mutants selected after exposure to aztreonam revealed a SNP in the *czcS* gene; this has previously been implicated in carbapenem resistance in a clinical isolate of this species. ^23^

Efflux is an important mechanism of bacterial multidrug resistance (MDR), and the inhibition of MDR pumps by efflux pump inhibitors (EPIs) could be a promising strategy to overcome MDR. Our previous work suggested that mutant selection experiments with a D408A mutant could be used in drug discovery programmes to identify substrates and/or competitive inhibitors of the main RND multidrug efflux pump in Gram-negative bacteria. ^9^ Exposure of a loss of function AcrB/MexB mutant to an AcrB /MexB substrate applies pressure to select for mutants with a wild type AcrB/MexB sequence and functional AcrAB-TolC efflux pump. It was hypothesized that there will be no evolutionary benefit to this reversion if these compounds were not pump substrates and so use of such a mutant in antibiotic exposure experiments could be used to reveal whether an antibiotic is a substrate of the pump.

Bacteria have evolved to have multiple mechanisms to evade the action of antibiotics, and our data indicates that when the lack of efflux via the major RND system in *E. coli, S*. Typhimurium and *K. pneumoniae* is important the only evolutionary path is to revert the mutation conferring loss of efflux function. When other routes of evolution to drug resistance are available, lack of efflux via this system is irrelevant. The same may be true for *P. aeruginosa*, but the lack of selection of wildtype *mexB* in our experiments may also be due to the difficulty in choosing a MexB substrate and experimental concentration with which to select mutants.

## Transparency declarations

None.

