## Supplemental tables 1-3 for "Mutation experiments with AcrB and MexB D408A mutants indicate the efflux liability of antibiotics"

**Supplementary Table 1. Primers used for constructing D408A mutation in *E. coli* and *K. pneumoniae acrB* genes.**

| Target | Primer name | Purpose | Sequence (5'–3') | Source |
|---|---|---|---|---|
| *E. coli acrB* | D408A F | Mutation cassette framing forward primer | CTTCCCGTCGGGTCTGAAAA | Wang-Kan *et al.*, 2021 |
| | D408A R | Mutation cassette framing reverse primer | AATCGGTTTCAGCATGGTGGC | |
| | pT2SKSM3 | Gibson assembly forward primer | ctaagcagggcggggcgtaaGTGCTCGCCATCGGCCTG | |
| | SM5pT2SK | Gibson assembly reverse primer | accgctgccactcttgagatTTCTTCCGCCATAACACGC | |
| *K. pneumoniae acrB* | D408A F | Mutation cassette framing forward | TTTACCCGTATGACACCACCC | This study |
| | D408A R | Mutation cassette framing reverse | ATAGTGATGGAGAACTGGCGATAGATG | |
| | pT2SKSM3 | Gibson assembly forward primer | gggcgtaaGGTGGATGCCGCCATCGT | |
| | SM5pT2SK | Gibson assembly reverse primer | cttgagatCCACGATGGCGGCATCCA | |
| | Kp AcrB D408Region F | Mutation check forward primer | TATTAAACTGGCTACCGG | |
| | Kp AcrB D408Region R | Mutation check reverse primer | AATGGGTTTCAGCATCGTGG | |

**Supplementary Table 2. Primers used to construct *P. aeruginosa mexB* (D408A) gene construct in pEX18 suicide vector**

| **pEX18 flanking primers** | **Primer sequence (5' – 3')*** |
|---|---|
| *pEX18-mexB* - forward | GACGGCCAGTGCCAAGCTTcatcatcggcaagacccg |
| *pEX18-mexB* - reverse | CAGGAAACAGCTATGACCATGATTACGAATTCgtcctcgtcggggag |
| **Mutagenic primers≠** | |
| *mexB* (D408A)- forward | GCCATCGGCTTGCTGGTGGACGcCGCCATCGTGGTGGTGGAGAA |
| *mexB* (D408A)- reverse | TTCTCCACCACCACGATGGCGTCGgCCACCAGCAAGCCGATGGC |
| **Check primers** | |
| *mexB* - forward | ACCTGTTCCTGCAGAACTTCC |
| *mexB* - reverse | ACGATGGTGATGGAGAACTGC |

*Bases in upper case overlapped with those on the pEX18 plasmid. ≠Red highlighted nucleotides are the substituted nucleotides.

**Supplementary Table 3. Real time PCR primers and cycling conditions**

| Species | Sequence |
|---|---|
| *E. coli/ K. pneumoniae acrB* 408A forward | 5' CATCGGCCTGTTGGTGGATAC 3' |
| *E. coli/ K. pneumoniae* | 5' CATCGGCCTGTTGGTGGATGA 3' |
| *E. coli acrB* reverse | 5' GATCGGCCCAGTCCTTCAAGGAAA 3' |
| *K. pneumoniae acrB* reverse | 5' CTCCAGTCTTTCAGCGAAACGAA 3' |
| *P. aeruginosa mexB* 408A forward | 5' CATCGGCTTGCTGGTGGACAC 3' |
| *P. aeruginosa* | 5' CATCGGCTTGCTGGTGGACGA 3' |
| *P. aeruginosa mexB* reverse | 5' CAGGGCTTGAGCATGATGAACG 3' |

**Cycling conditions**

*E. coli:* AcrB A allele 95°C (3 min), 95°C (30 sec) , 71°C (10 sec), 72°C (45 sec) (27 cycles). AcrB D allele 95°C (3 min), 95°C (30 sec) , 71°C (10 sec), 72°C (45 sec) (27 cycles).

*K. pneumoniae:* AcrB A allele 95°C (3 min), 95°C (30 sec) , 68°C (7 sec), 72°C (45 sec) (27 cycles). AcrB D allele 95°C (3 min), 95°C (30 sec) , 71°C (10 sec), 72°C (45 sec) (27 cycles).

*P. aeruginosa:* MexB A allele 95°C (3 min), 95°C (30 sec) , 63°C (10 sec), 72°C (45 sec) (27 cycles). MexB D allele 95°C (3 min), 95°C (30 sec) , 63°C (10 sec), 72°C (45 sec) (27 cycles).
